# Transitions in density, pressure, and effective temperature drive collective cell migration into confining environments

**DOI:** 10.1101/2023.04.10.536258

**Authors:** Wan-Jung Lin, Amit Pathak

## Abstract

Epithelial cell collectives migrate through tissue interfaces and crevices to orchestrate processes of development, tumor invasion, and wound healing. Naturally, traversal of cell collective through confining environments involves crowding due to the narrowing space, which seems tenuous given the conventional inverse relationship between cell density and migration. However, physical transitions required to overcome such epithelial densification for migration across confinements remain unclear. Here, in contiguous microchannels, we show that epithelial (MCF10A) monolayers accumulate higher cell density before entering narrower channels; however, overexpression of breast cancer oncogene +ErbB2 reduced this need for density accumulation across confinement. While wildtype MCF10A cells migrated faster in narrow channels, this confinement sensitivity reduced after +ErbB2 mutation or with constitutively-active RhoA. The migrating collective developed pressure differentials upon encountering microchannels, like fluid flow into narrowing spaces, and this pressure dropped with their continued migration. These transitions of pressure and density altered cell shapes and increased effective temperature, estimated by treating cells as granular thermodynamic system. While +RhoA cells and those in confined regions were effectively warmer, cancer-like +ErbB2 cells remained cooler. Epithelial reinforcement by metformin treatment increased density and temperature differentials across confinement, indicating that higher cell cohesion could reduce unjamming. Our results provide experimental evidence for previously proposed theories of inverse relationship between density and motility-related effective temperature. Indeed, we show across cell lines that confinement increases pressure and effective temperature, which enable migration by reducing density. This physical interpretation of collective cell migration as granular matter could advance our understanding of complex living systems.

## Introduction

As epithelia grow and migrate to sculpt organs, perform wound healing, or escape tumors, they must traverse physically heterogeneous environments of varying stiffness, topography, and confinement. Collectively migrating epithelial cells migrate faster on stiffer substrates due to higher levels of tractions forces associated with elevated actin-myosin activity (1, 2). When cells encounter an interface between soft and stiff regions, they undergo ‘durotaxis’ and quickly migrate into the stiffer side due to higher forces (3). On surfaces with gradient of stiffness, epithelial monolayers develop emergent durotaxis through force propagation across cell-cell junctions and seamlessly migrate from soft to stiff regions (4). Apart from stiffness, physical heterogeneities in the extracellular matrix (ECM) also arise from topography and confinement (5). According to previous work, both single and collective cells migrate faster in confined matrices, largely due to their alignment of forces and protrusion along a single axis (6-8). However, unlike the known durotaxis phenomenon for cell migration from soft to stiff regions (3, 4), it remains unclear how epithelial collectives migrate from open (unconfined) substrates into confined regions. Prior work has shown that collective cell migration is enhanced by cellular transformations of shape elongation, unjamming transition, and epithelial-mesenchymal transition (EMT) (9-11), all of which are promoted in confinement (12-14). Thus, upon encountering confined topographies, it is possible that epithelial collectives get arrested while undergoing such cellular transformation or they could seamlessly enter confinement analogous to durotaxis from soft to stiff environment; however, these possibilities remain unexplored.

Among the numerous physiological and pathological events related to collective cell behavior, epithelial-mesenchymal transition (EMT) is notable in its role in critical cellular processes such as embryogenesis and cancer invasion (15-18). Within epithelial clusters, cells are usually cuboidal, sedentary, have definite top-bottom polarity, and are in close contact with neighboring cells. During EMT, epithelial cells undergo a series of physiological changes, decreasing cell-cell junctions, elongating, and switching to front-rear polarity prior to migration. Mechanical properties of the environment, such as stiffness and topography, can also induce EMT. For example, stiff matrices triggers EMT in lung and breast epithelial cells without any growth factor addition (17, 18). On nano-grooved surfaces, switch-like enhancement of EMT is mediated by the Yes-Associated Protein (YAP) pathway (19). In breast cancer, the EMT-related oncogene ErbB2 (Her2/neu epidermal growth receptor family) was found to be overexpressed in advanced stage patients (20), and its overexpression enhances cell migration in confined matrices (21). Furthermore, confinement and stiffness could induce EMT in epithelial clusters independently (22-25). Since aligned matrix topographies trigger mechano-active responses, such as EMT, in epithelial cells (19, 22-25), collectively migrating epithelia could exploit such topography-sensitivities to enter confined matrices.

Moving beyond EMT, flocking and migratory behaviors of epithelial cells have also been described biophysically as analogues of solid-liquid phase transitions (26-30). Within cell clusters, where cells are usually constrained by neighboring cells, individual cell motions are coupled with their neighbors. At large distances and time scales, collective migration is analogous to particulate systems approaching a glass transition (26), wherein monolayer aging is analogous to solidification of glass particulates. Previous work has characterized shape and migratory variations of healthy and asthmatic human airway epithelial (HBEC; primary human bronchial epithelial cell) monolayers as “unjamming transition” (UJT). While baring some phenotypic resemblance with partial EMT (pEMT), the UJT triggered by compression is found to follow a distinct mode of cell migration (30-33). Existing theoretical and empirical models attribute these collective cellular transformations, via EMT, pEMT or UJT, to dynamic competition and cooperation between cell-cell junctions and intracellular forces (31, 32, 34). In this balancing act of forces and cohesion, RhoA (Ras homolog family member A), a member of small Rho GTPases, affects actomyosin contractility as well as cell-cell junctions (35, 36). By the differential levels of actomyosin contractility, leader-follower hierarchy within the migrating epithelial cluster is established and forward movement is ensured (16, 37, 38). Given RhoA itself could induce divergent downstream effects and thus opposing morphological changes (39), its activation in cell collectives could alter how they change shape and negotiate varying matrix topographies.

Given that cell migration is influenced by various properties of their microenvironment including confinement, stiffness, surface geometry and even external shear, most reports of mechanosensitive cell migration are cell-type specific or assay-specific (40, 41). Microchannels have been one of the popular models for confined cell migration (42, 43) since they allow easy manipulation and independent control of surface chemistry, material stiffness and physical constraints (5, 6, 22, 44). Emerging migratory modes inside microchannels versus channel conditions have been characterized either biochemically through signal pathways or physically by diffusivity and oscillation (22, 23, 45-48). Whereas researchers have previously studied collective cell migration in defined confinement, how cells collectively negotiate and move from unconfined to confined matrices remains unclear. We adopt a biophysical approach combining microchannels and mutant epithelial cells to study monolayer migration into confinement.

In this study, we use MCF10A (10A) human mammary epithelial cells and their derivative cell lines with constitutively-active RhoA (10A+RhoA) and overexpressed ErbB2 (10A+ErbB2) to form epithelial monolayers. We fabricate substrates with unconfined (open) region adjacent to confined (narrow or wide) channels to study monolayer migration into confinement. By analyzing migration phenotypes, individual cell shapes, and cell density, we found that monolayers develop differentials of density, pressure, and shape transitions between confined and unconfined regions. These migratory and shape transitions into confined regions reduce with EMT-like +ErbB2 mutant cells. Our findings indicate the collective cell migration into confinement is driven by cell-level physical changes of density, pressure, and effective temperature; yet these physical rules of migration must vary for cell types with healthy, cancerous or contractile traits.

## Results

### Epithelial monolayers undergo crowding, higher cell density, before entering confinement

To image and analyze how epithelial monolayers enter matrix regions of narrowing confinement, we fabricated PDMS substrates with unconfined region adjacent to microchannels (Figs. 1,S1) of widths 200μm and 50μm to capture wide and narrow confinements, respectively (Fig. 1). By tracking nuclei within monolayers entering microchannels, we calculated cell densities separately outside and inside the microchannels (Fig. 1c). Through live imaging over time, we observed that cell density is relatively higher in unconfined regions (outside) compared to inside microchannels, as shown in representative phase contrast and nuclei images in Figs. 1a,b. Cell density increases to higher levels when cells attempt to enter narrow channels, compared to wide (middle column in Figs. 1a,b), which continues to hold true even after cells migrate into the channels. We plotted average cell density over several samples overtime and found that cell density rises over the first 12 hours and stays higher outside the microchannels, compared to inside (Fig. 1d,e). Once the monolayers established stable flows inside microchannels, the cell density heatmaps revealed greater cell densities outside narrow microchannels, compared to those in the wide group (Fig. 1f).

**FIGURE 1.**
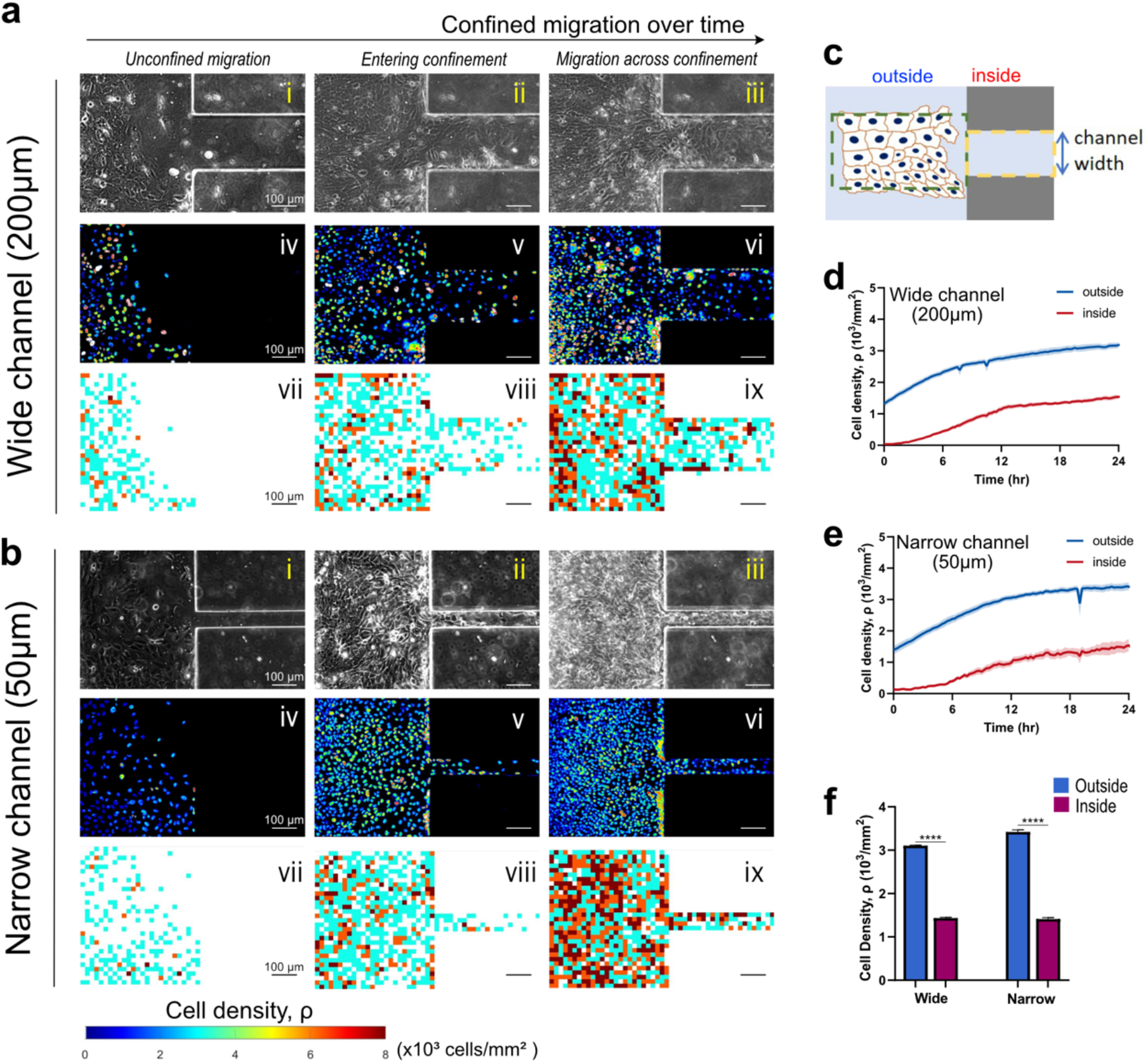
Epithelial monolayers build up cell density gradients to enter confining matrices. Representative snapshots of MCF10A epithelial monolayer entering **(a)** 200-µm (wide) or **(b)** 50-µm (narrow) microchannels shown at three key timepoints: (i, iv, vii) when monolayer reaches the channel wall; (ii, v, viii) intermediate process of monolayer attempting to enter the channel; (iii, vi, ix) final stage when the monolayer already establishes steady flow into the channel. (i-iii) Phase contrast images; (iv-vi) FIRE LUT rendering of GFP-labelled nuclei within the monolayer; (vii-ix) cell density heatmap. Scale bar: 100 µm. **(c)** A top-view schematic showing the regions of interests (ROI) “outside” and “inside” the channel. Cell density curves of WT monolayers entering **(d)** wide and **(e)** narrow microchannels over 24 hours, where blue and red curves represent average cell density outside and inside the microchannels, respectively. **(f)** Average cell density within the WT monolayers from *t* =18-24 hr is significantly different (*p* < 0.0001, assessed by two-way ANOVA) outside (blue) and inside (purple) microchannels in both wide and narrow groups (n≥8).

Thus, entry of collectively migrating epithelial cells into narrower confinement requires a rise in cell density. We wondered whether such accumulation of cell density near narrower confinement would also depend on cellular properties, specifically those regulating cohesion of epithelial population. Since cell monolayers maintained higher density outside (unconfined region) compared to inside microchannels in the wildtype 10A case (Fig. 1d-f), we calculated cell density (*Δρ*) difference between outside and inside microchannels over time and found it to be slightly higher in case of entry into narrow confinement (Fig. 2c). When WT monolayers entered the wide microchannels, the density difference *Δρ* peaked at 6 h after the monolayer reached the microchannel wall and dropped at 12 h, while the density difference of monolayers entering the narrow microchannel plateaued after 6 h (Fig. 1c).

**FIGURE 2.**
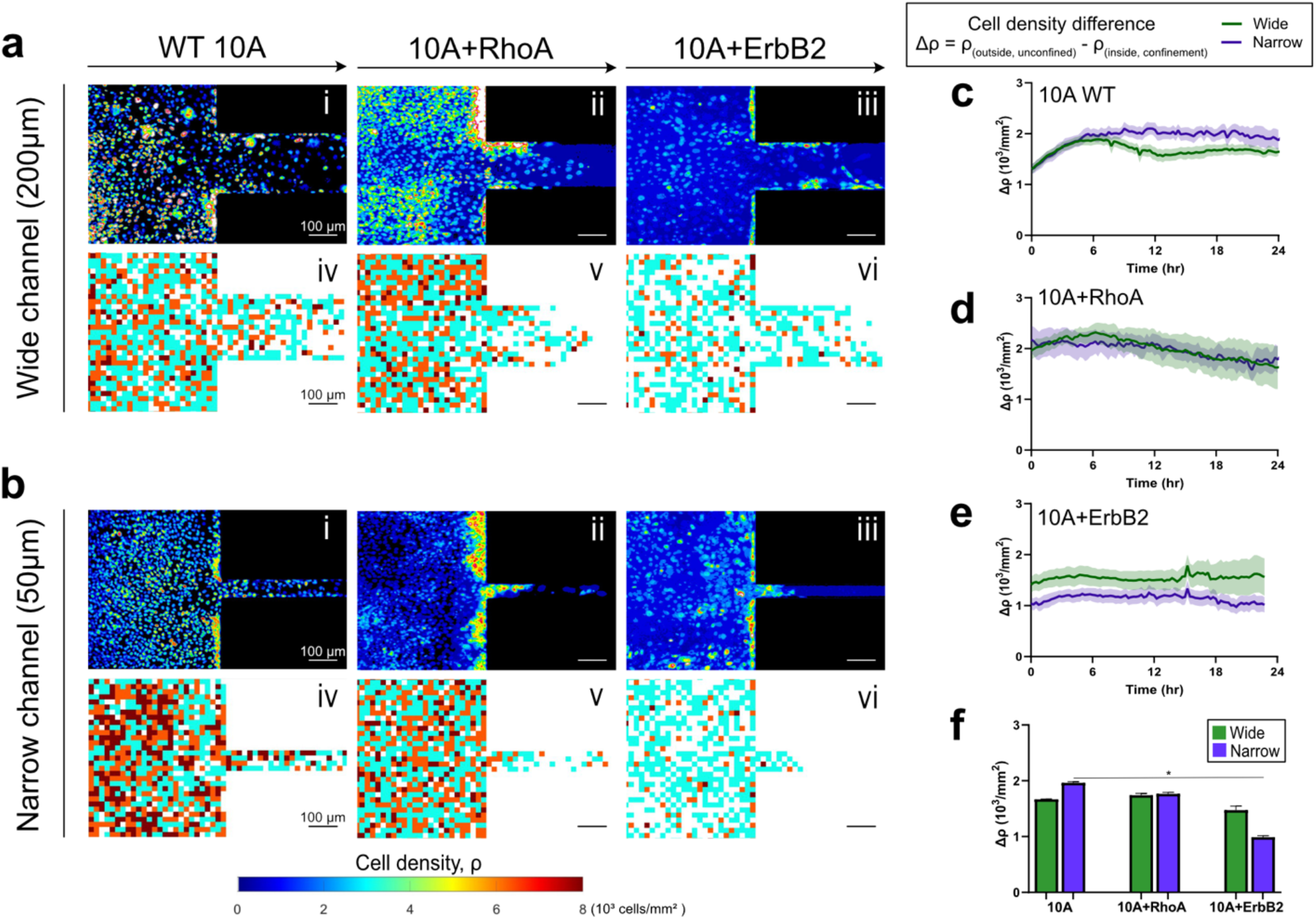
Dell density gradients across confinement vary with cell types. Cell density distribution after collective migration into **(a)** wide and **(b)** narrow channels for MCF10A wildtype, 10A+RhoA, and 10A+ErbB2 cell variants, with (i-iii) nuclear intensity and (iv-vi) cell density heatmaps. Temporal variation of difference in cell density between inside and outside the microchannels, *Δρ*, for **(c)** WT, **(d)** 10A+RhoA and **(e)** 10A+ErbB2 cells. **(b)** Cell density difference averaged from *t* = 18-24 hr. \**p* < 0.05, by two-way ANOVA. N≥4 ROIs.

Next, we used MCF10A cells with constitutively-active RhoA, which is known to enhance intracellular contractility and reinforce cell-cell adhesions (41, 49). We found that for the first 24 hours after monolayers reached channel walls, 10A+RhoA cell monolayers maintained similar cell density difference between outside and inside both narrow and wide channels (Fig. 2d). Although cells overexpressing RhoA also exhibited cell density accumulation outside of microchannels, they did not differentiate between narrow and wide channels. Furthermore, given cancer cells navigate confinement environments in tumor invasion and cancer metastasis, we used MCF10A cells with overexpressing oncogene ErbB2, which features in advanced stage breast cancer and relates to EMT (50, 51). Here, we did not use a different cancer cell line to maintain the same MCF10A cell line and observe different phenotype as one specific gene is altered. We found that 10A+ErbB2 cell monolayers maintained higher cell density difference *Δρ* across confinement when entering wide channels, compared to narrow, over 24hr duration of migration (Fig. 2e). For 10A+RhoA monolayers, the cell density difference necessary to enter the wide and the narrow channels were almost identical (Fig. 2d,f), whereas in 10A+ErbB2 monolayers, the cell density difference necessary to enter the narrow channels was smaller than that necessary to enter the wide channel (Fig. 2e,f). Thus, compared to 10A and 10A+RhoA cells, this lower density accumulation of 10A+ErbB2 cells (Fig. 2e,f), particularly while moving to narrow channels, indicates that cancer-like mutations could make it easier for cells to enter confined regions, without requiring density accumulation (52).

### Faster migration in confined regions and reduced confinement sensitivity by mutant cells

To assess spatiotemporal evolution of migration speed into confinement, we plotted average speed of individual cell migration tracks as monolayers of all three cell types entered narrow or wide microchannels (Fig. 3a-f). In these representative images, wildtype (10A) cells were faster than both mutant cells, with faster tracks going into the channels. Upon plotting root mean square (RMS) cell velocity (*υ*_*rms*_) over time for wildtype 10A cells, we found higher cell speed inside microchannels compared to the outside unconfined regions (Fig. 3d). For both +RhoA and +ErbB2 cells, migration velocity of cells outside and inside channels were similar (Figs. 3e,f). We calculated difference in *υ*_*rms*_ between outside and inside channel (*Δυ*_*rms*_) averaged over the last 6hrs of imaging, which indicates degree of sensitivity for confinement by a given cell type (Fig. 3g). We found that wildtype cells displayed the highest difference between confined and unconfined cell migration speed (Fig. 3d), which further increased when monolayers attempt to enter narrow channels. This velocity difference reduced for both mutants (+RhoA and +ErbB2), while cancer-like +ErbB2 cells moving into narrow channels exhibited the smallest velocity difference, indicating their lack of sensitivity for confinement.

**FIGURE 3.**
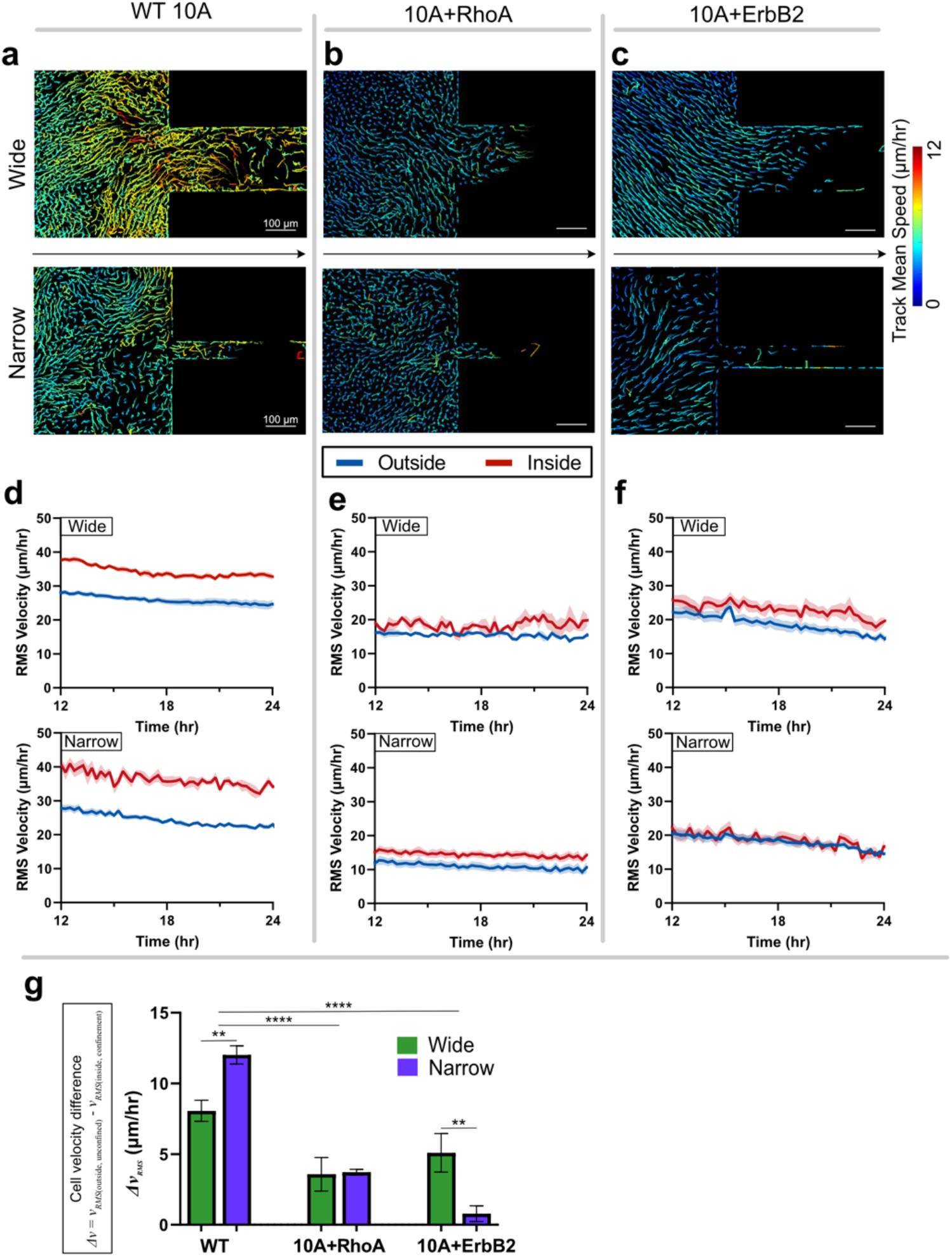
Cell velocity variations due to confinement and cell types. Migration tracks of cells within **(a)** MCF10A WT, **(b)** 10A+RhoA and **(c)** 10A+ErbB2 monolayers entering wide and narrow channels, with tracks color-coded according to the track mean speed. Root-mean-squared (RMS) velocities of **(d)** WT, **(e)** 10A+RhoA, and **(f)** 10A+ErbB2 cells outside (blue) and inside (red) channels. **(g)** Average difference in RMS velocity of cells outside and inside the channels. \*\**p* < 0.05, \*\*\*\**p* < 0.001

While epithelial cells collectively migrating in unconfined environments can stay together, their entry into confined regions requires some cell groups to be left behind. This spatial heterogeneity could cause migratory disorder at the leading edge, which could also vary with healthy and cancer-like cells. In tracking cells within monolayers, we observed that WT 10A cells outside of microchannels persistently maintained their direction of migration over time (Fig. S2a) regardless of narrow or wide channels. In comparison, directional persistence of 10A+RhoA cells reduced during 12-24hr time-window (Fig. S2b). With +ErbB2 mutation, migrating monolayers showed higher and temporally stable directional persistence than both WT and +RhoA variants (Figs. S2). We also calculated order parameter as cosine of angle between individual velocity vectors and direction of the channel, which largely resembled directional persistence trends (Fig. S2b). For +ErbB2 cells, while persistence was consistently higher in entering wide channels, order parameter for narrow channels increased over time, indicating a tendency of 10A+ErbB2 cells to expand regardless of confinement dimension.

### Migrating epithelial collectives develop pressure difference upon encountering confinement

In time-lapse videos, we observed that epithelial monolayers freely expanded and filled unconfined regions ahead of microchannels without actively entering immediately (Fig. 1). To further understand physical importance of such accumulation and reduced velocity of cells before entering confinement, we attempted to estimate pressure generated by the migrating monolayer, how it changes upon encountering sudden change in confinement, and how this pressure changes between inside and outside the channels. If the migrating collective followed the physical principles of simple fluid flow, the Bernoulli’s equation could be used to pressure from the known variations in velocity and density outside and inside confinement. Thus, we estimated the effective pressure difference experienced by migrating epithelial cells between outside and inside channels as *ΔP*_*eff*_ = *ρ*_*out*_*υ*_*out*_^2^ − *ρ*_*in*_*υ*_*in*_^2^ (Fig. 4). This pressure difference could be attributed to change in kinetic energy across narrowing spaces, previously estimated from cell migration speeds (53). By plotting overtime, we found that the pressure difference *ΔP*_*eff*_ in 10A wildtype cells started at a high level and approached zero after 12hr (Fig. 4), at which points cells began to enter confinement. Such physical description explains that the difference in pressure generated by epithelial monolayer between outside and inside confinement reaches an equilibrium before seamlessly entering confined environments. This difference in pressure across confinement *ΔP*_*eff*_ was lower in mutants cells compared to wildtype (Figs. 4b,c). Yet, resembling the pressure drop observed in wildtype cells, *ΔP*_*eff*_ for both +RhoA and +ErbB2 cells reduced to zero as cells migrated into confinement. Additionally, the wave-like trend of the *ΔP*_*eff*_ curves (Fig. 4) could be associated with intrinsic oscillations within the epithelial monolayers (48). The velocity difference between outside and inside channels (Fig. 3) suggests a physical segregation within the monolayers. Thus, although the level of pressure difference *ΔP*_*eff*_ varies with cells’ intrinsic properties, the physical requirement of reduction in pressure for continued collective cell migration into confinement holds true across cell types. Considering these trends in effective pressure, we speculate that cell shapes and their effective temperature could also undergo transitions to enter confining environments.

**FIGURE 4.**
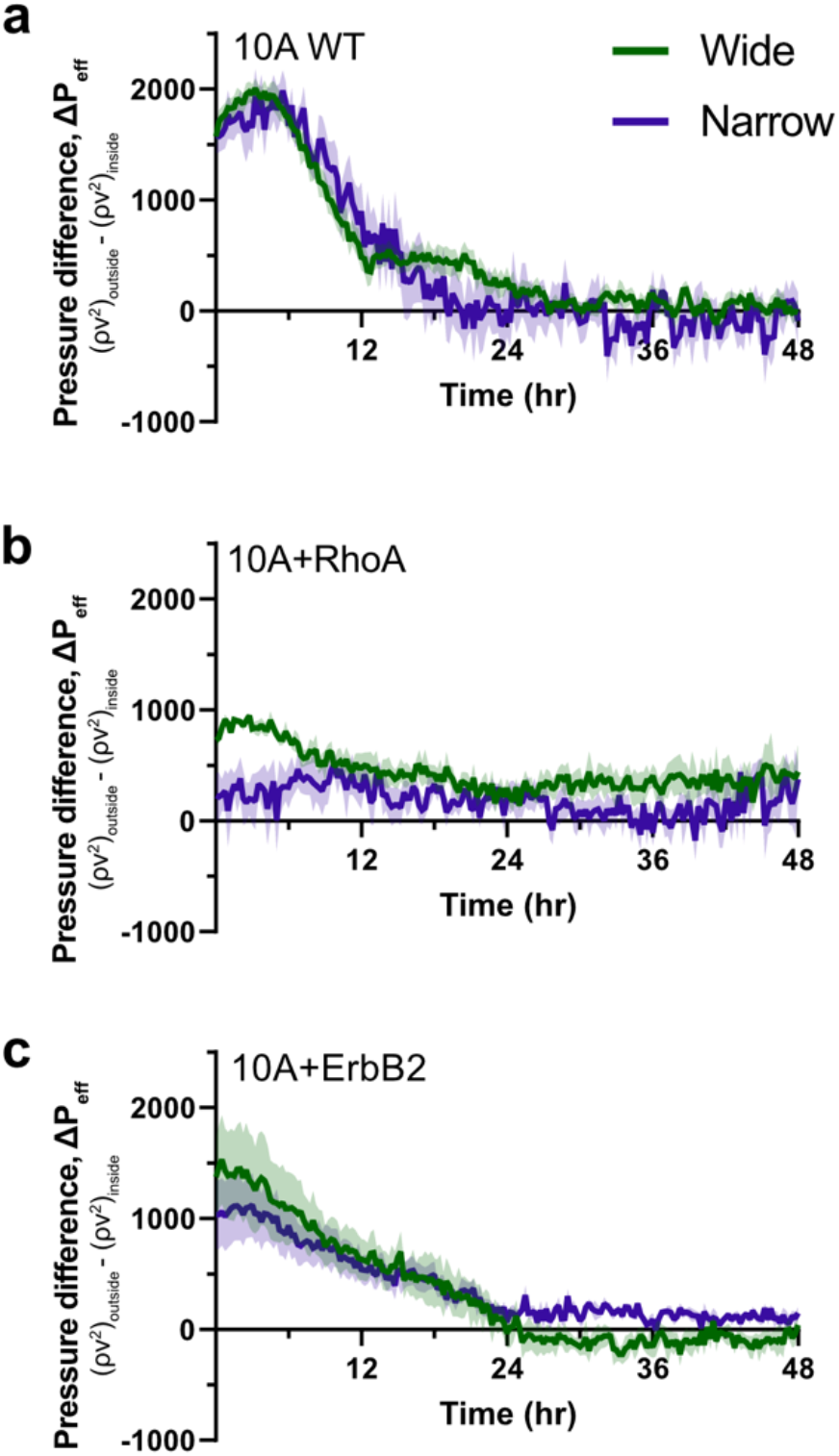
Effective pressure difference between inside and outside confinement. Temporal variation of effective pressure difference *ΔP*_*eff*_ = *ρ*_*out*_*υ*_*out*_^2^− *ρ*_*in*_*υ*_*in*_^2^ for **(a)** MCF10A WT, **(b)** 10A+RhoA and **(c)** 10A+ErbB2 monolayers entering wide (green) and narrow (purple) channels in 48 hours.

### Cells undergo mixed morphological changes of jamming, unjamming, and elongation to enter confinement

Since epithelial monolayers develop gradients of density and pressure (Figs. 1-4) across confined and unconfined regions, we next wondered whether individual cell shapes change during these transitions of encountering and entering confinement. In previous studies, cell elongation has been calculated as aspect ratio (AR), and values of shape index (SI; perimeter divided by square-root of area) ≥3.81 indicate unjamming (11). Furthermore, SI and AR of cells have been used to described different states of UJT and pEMT (32) such that low values of both SI and AR indicate jamming, high values of both SI and AR indicate UJT, and low AR combined with high SI indicate pEMT. To segment individual cell outlines with epithelial monolayers, we performed immunostaining for p120, which localizes on adherens junctions (Fig. 5a). Through custom cell segmentation tools, we visualized individual cells within migrating monolayers by color-coding for their AR (Fig. 5b), SI (Fig. 5c), and combined AR and SI traits (Fig. 5d). From spatial shape distributions, we observed that majority of cells had 1≤AR≤3 and 4≤SI≤5 (Fig. 5b, c, f, g), which suggests that most cells underwent moderate elongation yet developed UJT-like high shape index. In WT 10A and +RhoA monolayers (Fig. 5b, c), cells with very high AR (≥4) or highly unjammed SI (≥6) shape index, both in red hues, were found at the entry of microchannels, indicating their need to undergo larger than usual shape transitions to enter confinement. In comparison, 10A+ErbB2 cells had more homogeneous distribution of AR and SI, with very few cells showing large shape changes, indicating their less requirement for dramatic shape transitions to enter confinement. We plotted average metrics for cell morphology across samples and found that cell area for WT and +RhoA cells was lower than +ErbB2 (Fig. 5e), further indicating the need for non-cancer cells to get squeezed to enter confinement. While average AR of WT and +RhoA cells was higher than ErbB2, differences in SI across cell lines were statistically insignificant when averaged over the whole population (Figs. 5f, g.)

**FIGURE 5.**
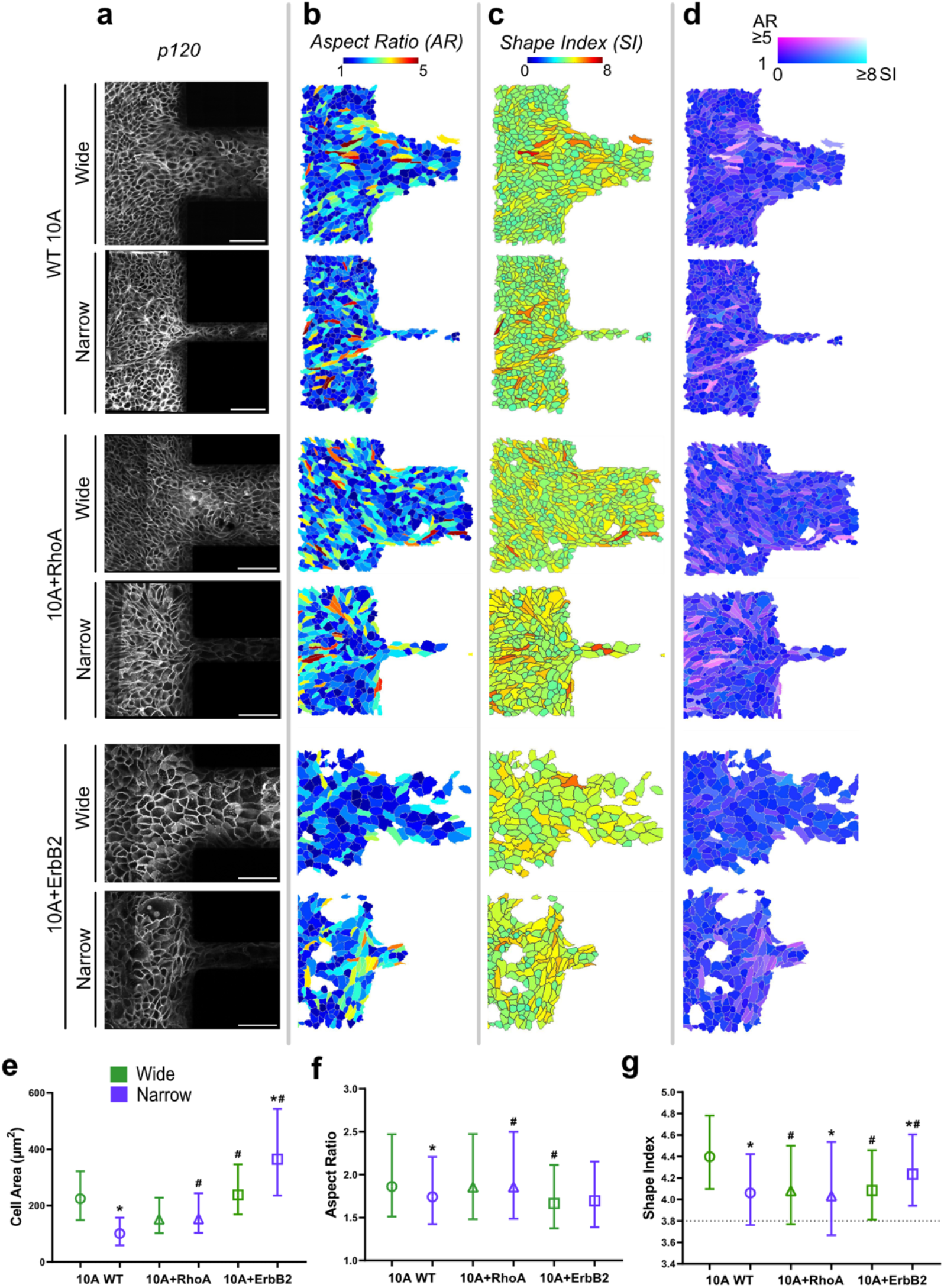
Morphologies of cells in epithelial monolayers entering confinement. For MCF10A WT, 10A+RhoA and 10A+ErbB2 monolayers entering the wide and narrow channels, **(a)** immunostaining of cell-cell junctions (P120), **(b)** aspect ratio (AR), **(c)** shape index (SI), and **(d)** combined representation AR and SI according to the indicated colormap. Comparison of averaged (e) cell area, (f) aspect ratio, and (g) shape index across cell type and confinement variations, where the markers and bars represent median and interquartile range, respectively. Kruskal-Wallis test is used here for pairwise comparison between groups, *p<0.05 compared to wide channels. #p<0.05 compared to WT. N≥1700 cells from at least 4 ROIs.

We also visualized morphological state of each cell as combined effect of AR and SI (Fig. 5d) to spot cells with relatively lower unjamming (SI) yet higher elongation (AR). This visualization allows assessment whether cells get jammed despite getting elongated while entering confinement. Indeed, in WT 10A and 10A+RhoA monolayers, many cells showed magenta hues (Fig. 5d) of higher AR and lower SI, which indicates that cells undergo some jamming despite shape elongation required to enter confinement. These jammed-elongated cells (darker magenta) seemed to be squeezed ahead of channel entry while unjammed-elongated cells (lighter magenta) migrated into channels. For +ErBb2 cells, the jammed-elongated (dark magenta) phenotype was minimal, and most cells showed unjamming (blue), particularly in wide channels. When migrating into narrow channels, +ErbB2 showed some jamming behavior (magenta cells), but this phenotype of was much less prevalent compared to WT or +RhoA cells.

### Pharmacological epithelial reinforcement of cancer-like mutant cells suppresses unjamming

According to our measurements, the cancer-like 10A+ErbB2 entered confinement with smaller gradients in cell density (Fig. 2) and migration speed (Fig. 3) while retaining lower pressure (Fig. 4) and homogenous shape distributions (Fig. 5). Since ErbB2 is an EMT-related oncogene (20), it is possible that +ErbB2 monolayers are less cohesive than WT 10A and +RhoA counterparts, which allows them to move more seamlessly into confinement. Thus, we wondered whether reinforcement of epithelial characteristics in 10A+ErbB2 cells could reverse some of these phenotypes towards those exhibited by the cohesive cells (WT or +RhoA) in our study. To this end, we treated 10A+ErbB2 with small doses of Metformin, a diabetes drug that interferes with cell cycle progression, enhances cell-cell adhesions, and alters focal adhesion turnover rate (54). The outside-inside cell density difference *Δρ* of metformin-treated +ErbB2 cells for wide channels was higher in the beginning (Fig. 6b), resembling the +ErbB2 cells (Fig. 2e). However, over time the metformin-treated cells closed this wide-narrow gap in cell density and resembled +RhoA cells (Figs. 6b, c). The difference in cell migration speed between wide and narrow channels observed for 10A+ErbB2 cells vanished after metformin treatment, resembling the +RhoA cells (Figs. 6e). While the directional persistence of untreated 10A+ErbB2 cells was higher in wide channel, metformin treatment reversed this trend and yielded higher persistence in narrow channels (Fig. 6d), which also resembles +RhoA cells. We also performed morphometric analyses and found that metformin treatment introduced some cells with high AR and SI (AR≥4 and SI≥6, visualized in red hues Figs. 6g, h), as was earlier observed for WT and +RhoA cell but not for untreated +ErbB2 cells (Figs. 5f, g). In combined visualization of AR and SI, some cells exhibited high AR with low SI (Fig. 6i), which is the so called jammed-elongated cell shape observed earlier (Fig. 5) in cohesive cell monolayers without the +ErbB2 mutation. By plotting averaged morphology metrics across samples, we found significant reduction in cell area (Fig. 6j), indicative of cells getting squeezed, as well as slight reduction in aspect ratio and shape index (Fig. 6k,l), indicative of jamming signatures. Taken together, metformin treatment shifted both migratory behavior and morphological composition of +ErbB2 cells towards the relatively more cohesive WT and +RhoA cells.

**FIGURE 6.**
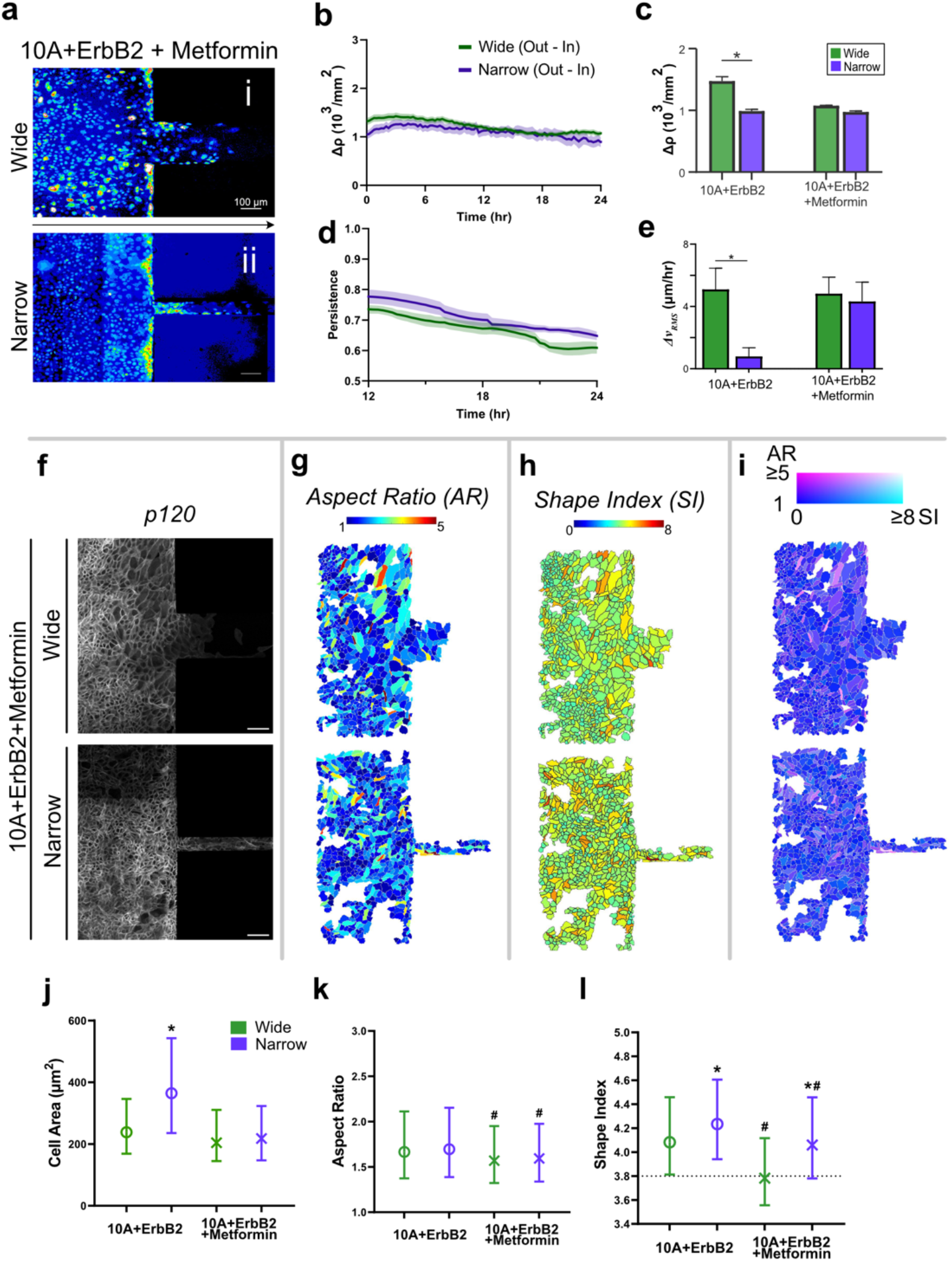
Cell migration and morphologies after treatment of 10A+ErbB2 cells with Metformin. **(a)** Nuclear intensity within 10A+ErbB2+Metformin monolayers entering (i) wide and (ii) narrow channels. **(b)** Temporal variation of cell density difference *Δρ* between outside and inside channels, **(c)** averaged *Δρ*, **(d)** temporal persistence, and **(e)** averaged difference in RMA velocity *Δv*_*RMS*_ between outside and inside channels for 10A+ErbB2+Metformin cells. \**p* < 0.05. **(f)** P120 expression from immunostained images, **(g)** aspect ratio, **(h)** shape index and **(i)** combined colormap of +ErbB2+Metformin monolayers. Comparison of averaged **(j)** cell area, **(k)** aspect ratio and **(l)** shape index of 10A+ErbB2 cells with and without Metformin treatment, where markers and bars represent median and the interquartile range, respectively. Kruskal-Wallis test is used for pairwise comparison between groups. *p<0.05 compared to wide channels. #p<0.05 compared to ErbB2 untreated control. N≥3400 cells from at least 8 ROIs.

### Inversely correlated cell density and effective temperature in confined migration

Epithelial cells accumulate higher density (Fig. 2) and generate wide distribution of cell shapes (Fig. 5) as they enter confinement, with varied combinations of jamming, unjamming and pEMT phenotypes for different cell types. To better understand these cell shape transitions in physical terms, we described epithelial monolayers as a thermodynamic system of granular matter, as done previously (10). We used the *k*-gamma distribution *PDF*(*x*;*k*)= *k*^*k*^*x*^*k*-1^*e*^−*kx*^/*Γ*(*k*) to fit the histograms of rescaled aspect ratios 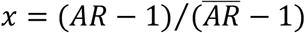) (31). From this fitting, we calculated *k* and 95% confidence intervals (CI) and effective temperature 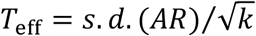) of monolayers across wide and narrow channels, as listed in Table 1. According to these calculations, 10A+RhoA reached the highest effective temperature *T*_*eff*_, with slight reduction in narrow channels, which is consistent with their propensity to jam and high heterogeneity in cell shape (AR and SI) measured earlier (Fig. 5). By contrast, *T*_*eff*_ of 10A+ErbB2 cells was lower, with slightly higher temperature in narrow channels, which is consistent with earlier observation of ability of enter confinement more easily with minimal density gradients and jamming. Thus, +RhoA and +ErbB2 respond to degree of confinement in opposing manner, with +RhoA warmer in wide channels and +ErbB2 warmer in narrow channels. For WT cells, microchannels caused limited jamming and slightly cooled the monolayers.

**Table 1.**

| Cell lines | Channel width | Cell number | k | 95% CI | $T_{eff}$ |
| --- | --- | --- | --- | --- | --- |
| **10A WT** | 200 $\mu\text{m}$ | 1704 | 1.91 | [1.79, 2.02] | 0.6054 |
| | 50 $\mu\text{m}$ | 4669 | 1.82 | [1.75, 1.89] | 0.5467 |
| **10A+RhoA** | 200 $\mu\text{m}$ | 6570 | 1.71 | [1.66, 1.77] | 0.6834 |
| | 50 $\mu\text{m}$ | 4595 | 1.74 | [1.67, 1.81] | 0.6784 |
| **10A+ErbB2** | 200 $\mu\text{m}$ | 6719 | 1.79 | [1.73, 1.84] | 0.5011 |
| | 50 $\mu\text{m}$ | 3499 | 1.74 | [1.66, 1.81] | 0.5737 |
| **10A+ErbB2**<br>**+Metformin** | 200 $\mu\text{m}$ | 5068 | 1.78 | [1.72, 1.85] | 0.4615 |
| | 50 $\mu\text{m}$ | 3496 | 1.78 | [1.71, 1.86] | 0.4601 |

In an expanding monolayer, bulk stresses are spatially constant and reach homeostatic stress (55) proportional to log(*ρ*), where *ρ* is local cell density. We suppose that the expanding monolayer experiences a normal force which is equal to the sum over the normal projection of its homeostatic stress on the channel wall. By assuming the monolayer is large enough with uniform cell density locally, we consider log(*ρ*) as a measure of internal stress that monolayer develops upon hitting channel walls and at the channel entrance, while *T*_*eff*_ could be a measure of cell motility. Building on a previous postulation that multicellular collectives evolve in the close vicinity of jamming state with fluid-like features (56), we plotted phase diagrams as the natural logarithm of cell densities, log(*ρ*), versus *T*_*eff*_ (Fig. 7a, b). When entering wide channels (Fig. 7a), linear regression of log(*ρ*) versus *T*_*eff*_ plots indicates that temperature (*T*_*eff*_) varies somewhat independently of the density-related stress log(*ρ*), with +RhoA and +ErbB2 cells showing higher and lower stresses, respectively, compared to WT. The low R^2^ in case of wider channels suggests change in stress is not necessary to raise *T*_*eff*_ and enhance migration into wide confinement. By contrast, in narrow channels (Fig. 7b), density and temperature are negatively correlated, indicating that loss of density-related stress, log(*ρ*), is necessary to reduce temperature *T*_*eff*_ and thus enhances motility.

**FIGURE 7.**
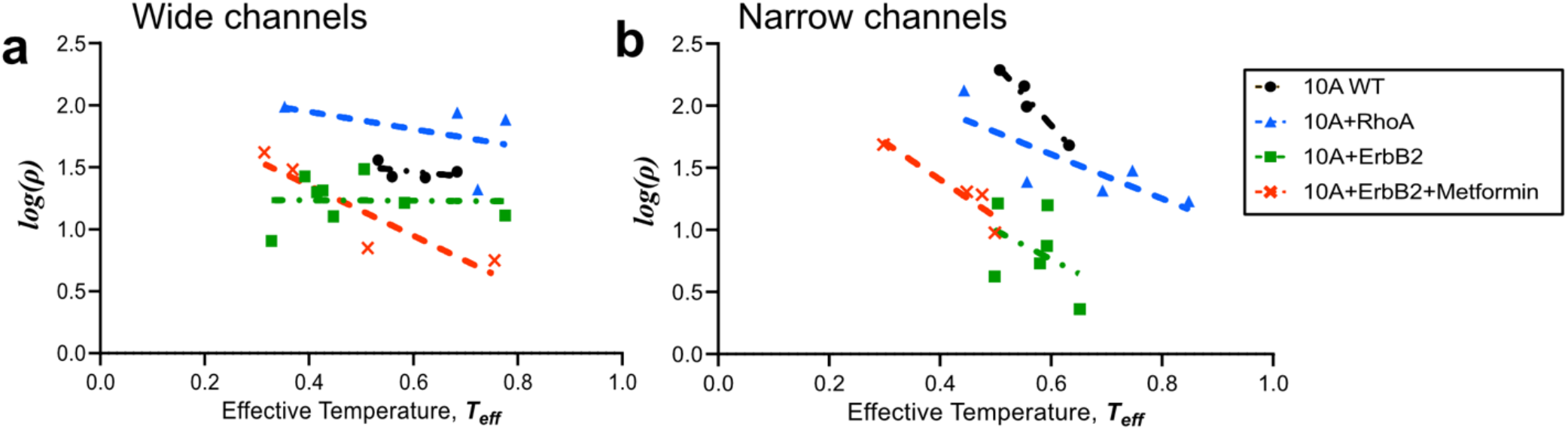
Inverse relationship between density and effective temperature depends on matrix confinement. Scatter plots of effective temperature *T*_*eff*_ versus the natural logarithm of local cell densities *log(ρ)* of monolayers entering the **(a)** wide and **(b)** narrow channels, along with the linear regression of each cell lines. N≥1700 cells analyzed from at least 4 ROIs for each condition.

## Discussion

Seventy years ago, Abercrombie & Heaysman posited an inverse relationship between cell density and migration for chick fibroblasts growing in a dish (57). This rise in density occurs over days, as regulated by cell proliferation, could cause the cell population to ‘jam’, although not characterized as such at the time. Over the past decade, cell collectives have been physically described as jamming of deformable bodies at high density, while unjamming requires rise in internal forces and shape transitions (11, 58). In theory, rise in temperature in thermally sensitive collective systems could achieve similar unjamming; whether caused by internal forces or temperature, unjamming has been associated with decreasing density (59). In epithelial monolayers, cell shape transitions have been used to calculate effective temperature by treating cells as a granular thermodynamic system (10). Thus, we wondered whether the inverse relationship between density and temperature, as theorized earlier (59), holds true in collective cell migration. Since the process of epithelial collectives approaching confinement would naturally incur crowding due to the narrowing space, cell density is expected to rise quickly within hours, without proliferation that needs days. Thus, our system of open-to-confined transmigration of collective cells allowed a unique opportunity to correlate cell shape-dependent effective temperature with density across cell lines.

In our system of epithelia attempting to enter microchannels, some cells hit the channel walls and revert their direction while others enter the microchannel. This mixture of tissue collisions and free expansion is similar to a recent tri-tissue collision experiment (60), except with the complication of forced and rapid changes in cell crowding and density. Indeed, after part of the monolayer hits the channel wall, the cells that do not enter the microchannel cause crowding outside, which creates a density gradient between inside and outside of the microchannel. When +RhoA cells approach confinement, their rise in density is similar in wide and narrow channels (Fig. 2). Since RhoA is associated with both cell contractility and cell-cell adhesions, it is possible that even the confinement presented by wide channels is rapidly sensed by this cell population due to their strong cohesivity. In wide type 10A cells, narrow channels are associated with slightly higher crowding than wide channels. By contrast, +ErbB2 cells show lower crowding ahead of narrower channels (Fig. 2), which could be attributed to their reduced contact inhibition, relation with EMT, and the cancer-like ability to crawl on top of each other (52). As a result, +ErbB2 cells do not require standard crowding and higher density accumulation ahead of narrower channels. Due to the ability of 10A+ErbB2 cells to crawl one another, they appeared to form 3D-like structures upon entering microchannel, while 10A+RhoA monolayers generally maintain their 2D monolayer phenotype. This contrast in macroscale structural differences could be attributed to stronger cell-cell contractility and cell-ECM adhesions downstream of RhoA (41, 61) and contact inhibition phenotype of +ErbB2. Since Wnt pathway is associated with intercellular separation and it involves the ErbB2 oncogene, we inhibited Wnt signaling using metformin as an attempt to restore contact inhibition in +ErbB2 cells (54, 62).

Through pressure and effective temperature calculations, we describe entry into microchannels as a way for the cell populations to minimize their pressure differential between the distinct confined and unconfined spaces. Due to this rise in density, when WT or +RhoA cells approach confinement, their crowding cools the system, calculated through cell shape-dependent effective temperature. Subsequently, some cells get squeezed and elongated, which warms the system as cell in free expanding portion of the monolayer enter channels (Fig. 5). Importantly, such physical coupling of crowding, temperature, and pressure is highly dependent on cells’ intrinsic properties. For instance, WT cells develop higher pressure upon encountering change in matrix space, while both mutants (RhoA, ErbB2) approach confinement with relatively lower pressures (Fig. 4). Yet, in all cases, pressure differentials between outside and inside the channels equilibrate over time as cells migrate into confinement. According to our physical interpretations, despite dissimilar cell types, collective cell migration seems to follow some basic relationships between density, shape-dependent temperature, and pressure minimization through migration into confinement. Our system of confinement gradient could also enable further studies of densification-driven migration, which is otherwise not feasible in situations with free boundary and growth-dependent densification of cell populations. Overall, these insights and physical analyses of cell migration could enable better understanding of how cell populations aggregate yet migrate in tumor invasion, wound healing, and embryogenesis.

## Materials & Methods

### Fabrication of PDMS substrates with microchannels

Microchannels of polydimethylsiloxane (PDMS, sylgard 184; Dow Corning, Midland, MI) were fabricated following the standard procedures of soft lithography, as depicted in Figure S1. Spin-coated SU-8 2050 (Kayaku Advanced Materials, Westborough, MA) photoresist was exposed and developed to yield grooves in sets of microchannels, each of widths 50μm (narrow) and 200μm (wide), spaced 700μm apart, and 60-70μm height. For easy release of cured PDMS, the master SU-8 2050 photoresist was treated with trichloro(1H,1H,2H,2H-perfluorooctyl)silane (448931, Sigma-Aldrich, Saint Louis, MO) in a desiccator for overnight. The monomers and crosslinkers of PDMS were mixed in a 10:1 ratio and degassed until clear. The mixed PDMS was spin-coated on the SU-8 master and incubated at 70°C for at least 2 hours and let sit at room temperature overnight; after fully cured, the PDMS was peeled from the master. The cured PDMS microchannels were cut into pieces and sealed to glass-bottom 12-well plates (P12-1.5H-N, Cellvis, Mountain View, CA) using ethanol. For the channel tops, thin strips of PDMS were plasma-bonded to the bottom layer and treated for one additional minute with air plasma to ensure the hydrophilicity of microchannel surfaces. The microchannels were then immersed in phosphate buffered saline (PBS; Thermo Fisher, Waltham, MA) and UV-sterilized for 1 h and coated with 0.05 mg/mL rat tail collagen I (sc-136157, Santa Cruz Biotechnology, Dallas, TX) at 4°C overnight.

### Cell culture

Human mammary gland wildtype (WT) MCF10A (10A) epithelial cells with green fluorescence protein-labelled nuclei, MCF10A cell line with constitutively active RhoA (10A+RhoA; courtesy of Gregory Longmore, Washington University in St. Louis), and MCF10A cell line with overexpressing ErbB2 (10A+ErbB2; courtesy of Dihua Yu, The University of Texas MD Anderson Cancer Center) mutant cell lines were cultured using Dulbecco’s Modified Eagle Medium/Nutrient Mixture F-12 (DMEM/F12; Invitrogen, Waltham, MA), supplemented with 5% horse serum (Hyclone heat-inactivated, Cytiva, Marlborough, MA), 20 ng/mL epidermal growth factor (EGF; Miltenyi Biotec, Gaithersburg, MD), 0.5 mg/mL hydrocortisone (H0888, MilliporeSigma), 100 ng/mL cholera toxin (C8052, Sigma-Aldrich), 10 μg/mL insulin (I6634, Sigma-Aldrich) and 100 μg/mL proprietary antimicrobial (Normocin; InvivoGen, San Diego, CA). Cells were cultured at 90-95% confluency, every three to four days. To seed epithelial monolayers, cell suspensions were prepared at 10^7^ cells/mL and 10 μL volume (10^5^ cells) droplets were used to grow monolayers in circular stencils of 3.7 mm diameter. These stencils were removed 1 h after seeding, and the wells were refilled with media. Epithelial monolayers formed within a few hours after seeding and migrating edges were 1-2 mm away from the channel entrance. A subset of 10A+ErbB2 monolayers were treated with 2 mM metformin (317240, Sigma-Aldrich) 2 h prior to live imaging, which was used to suppress mesenchymal phenotype triggered by overexpressing ErbB2.

### Live imaging and analysis of collective cell migration

After epithelial cells formed monolayers on PDMS substrates and approached microchannel entrances, live time-lapse imaging was performed. 10A+RhoA and 10A+ErbB2 cell lines were live stained with NucBlue (1:300; ThermoFisher, Waltham, MA) for 8 min and verapamil (1:5000-6000; Thermo Fisher) was used for preventing dye efflux from the nuclei. Images were acquired every 15 min for 36-72 h. Monolayer dynamics were analyzed using the open-source image processing platform Fiji (63) plugin TrackMate (64), which allowed us to track individual cell migration within the monolayers using the fluorescent nuclei as focal points. The “outside” region was defined as the 450 to 550 μm square region immediately in front of the microchannel entrance, and the “inside” region as the 450 to 550 μm square region immediately inside of the channel. The initial time point (t_0_) was defined as the instant the monolayers reached the channel walls (spanning length ≥ 25% in the field of view). TrackMate generated color-coded tracks according to the mean speeds and persistence throughout the duration of the experiment. The order parameter was defined as the projection of nucleus velocity along the channel direction. The persistence was defined as the total distance a nucleus migrated divided by the net displacement in 2 hours. The cell density heatmaps were obtained using nucleus positions given by TrackMate and custom MATLAB codes.

### Immunostaining and analyzing cell morphology

Cells were fixed for immunostaining 1-2 days after live imaging. Hoechst 33258 (1:50; Thermo Fisher), primary antibody of P120 (1:600, (D7S2M) XP; Cell Signaling Tech., Danvers, MA) and phalloidin (1:200, R415, Invitrogen) were used for labelling nuclei, P120 (δ-catenin) and F-actin. Images of P120 immunostaining (or those overlaid with F-actin images as necessary) were first used to determine cell outlines. The Python program SeedWaterSegmenter (v0.5.7.1, SWS) (65) was used for segmentation. Cell areas, aspect ratios and shape indices were calculated accordingly. The heatmaps of shape indices and aspect ratios were generated using a customized MATLAB code.

## Acknowledgements

This work was supported by the NIH/NIGMS MIRA (R35GM128764) grant to AP. Microscopy with live nuclear tracking was carried out at the Washington University Center for Cellular Imaging (WUCCI).

## Author Contributions

AP and WL conceived the project. WL performed experiments and analyzed data. WL and AP designed experiments, interpreted findings, made figures, and wrote the manuscript. AP edited the manuscript, supervised the project, and acquired funding.

## Supplementary Figures

**Figure S1.**
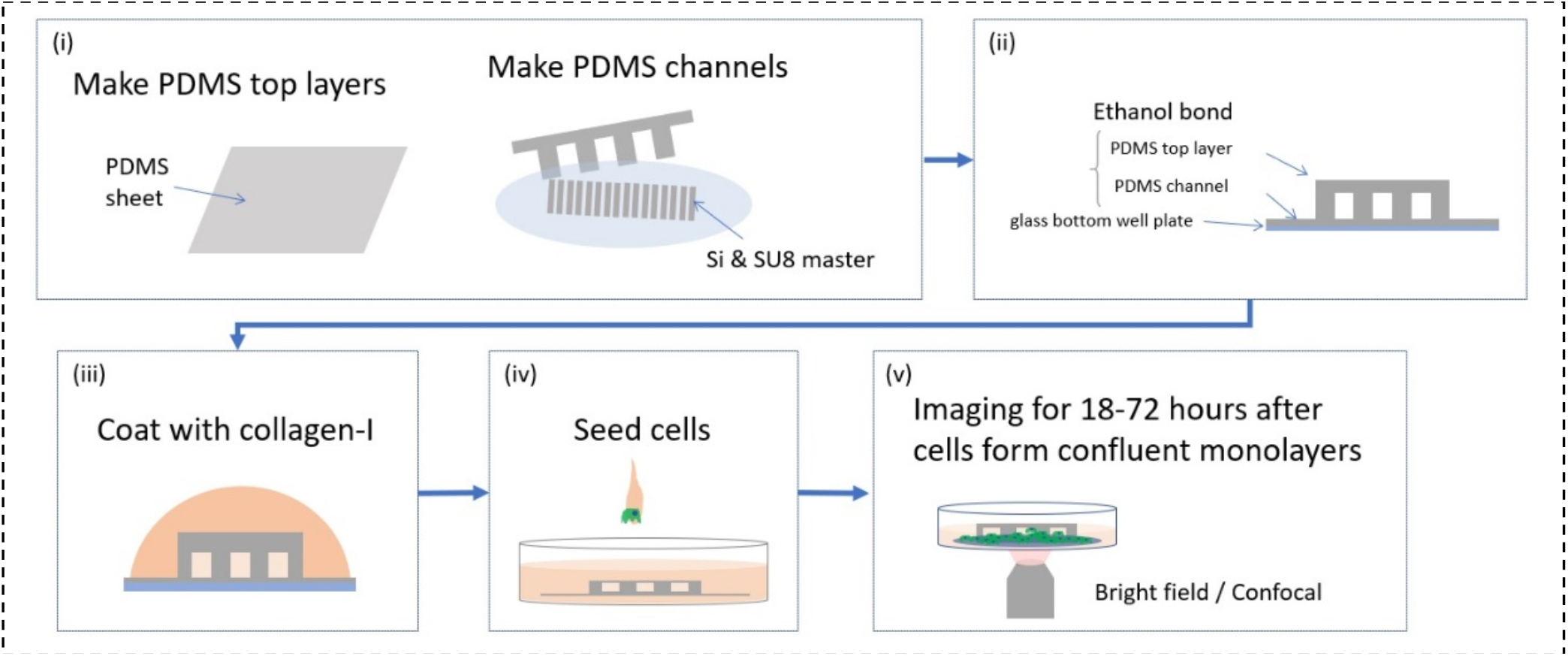
Experiment steps. (i) PDMS molding. (ii) Ethanol bonding of PDMS channels. (iii) Coating all surfaces with collagen type 1. (iv) Seeding cells outside the channels. (v) Live timelapse imaging with epifluorescence or confocal microscopy.

**Figure S2.**
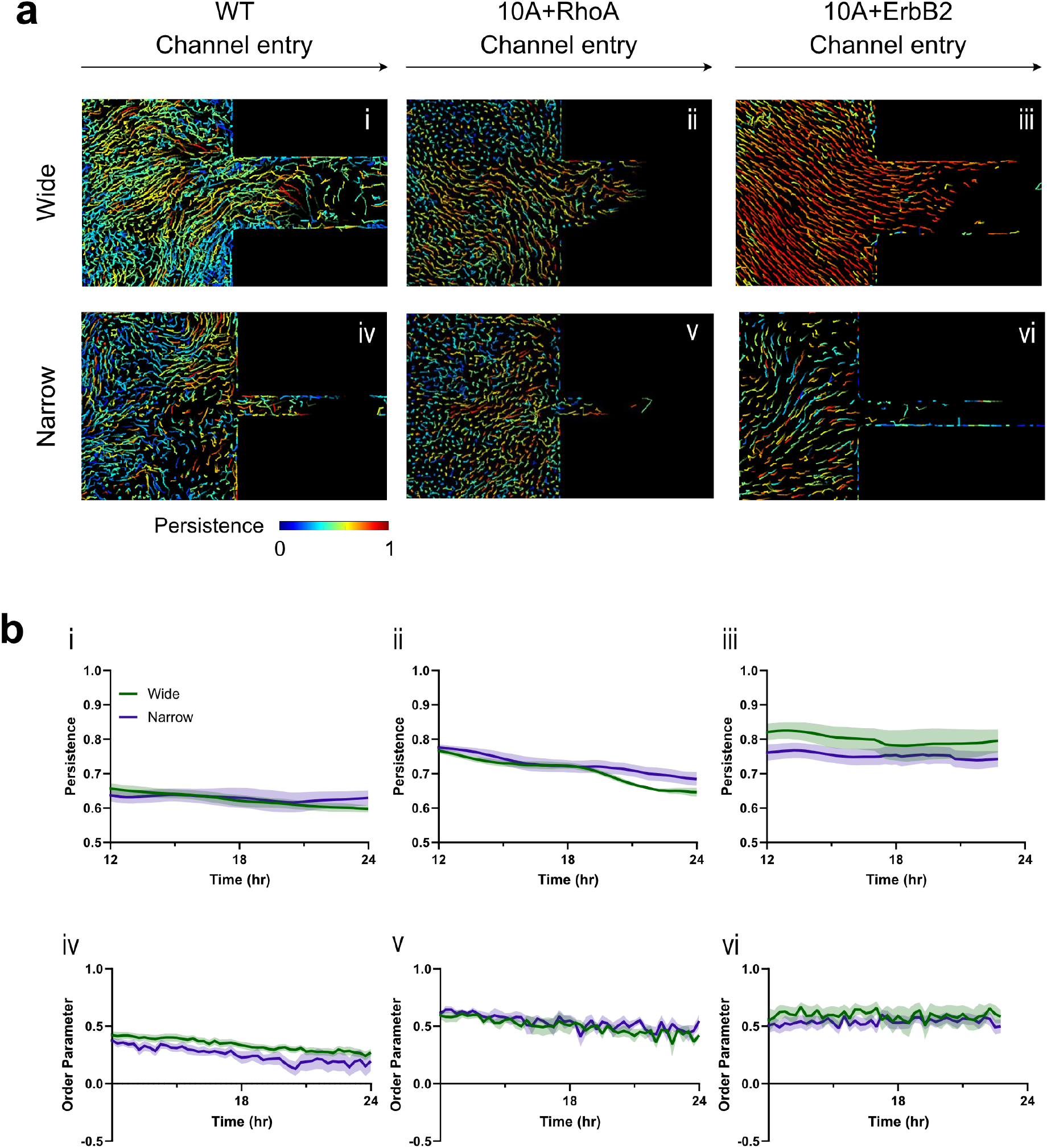
Persistence and order parameter within the monolayers across cell lines. (a) The snapshots of cell tracks within the monolayers color-coded according to track-wise persistence. (b) Curves of persistence and order parameter within the monolayers across cell lines. (i) WT_wide: n = 17; (ii) 10A+RhoA_wide: n = 4; 10A+ErbB2_wide: n = 7; (iv) WT_narrow: n = 8; (v) 10A+RhoA_narrow: n = 6; (vi) 10A+ErbB2_narrow: n=18.

## References

1. M. R. Ng, A. Besser, G. Danuser, J. S. Brugge, Substrate stiffness regulates cadherin-dependent collective migration through myosin-II contractility. J Cell Biol 199, 545–563(2012).

2. X. Trepat et al., Physical forces during collective cell migration. Nature Physics 5, 426–430(2009).

3. C.-M. Lo, H.-B. Wang, M. Dembo, Y.-l. Wang, Cell Movement Is Guided by the Rigidity of the Substrate. Biophysical Journal 79, 144–152(2000).

4. R. Sunyer et al., Collective cell durotaxis emerges from long-range intercellular force transmission. Science 353, 1157–1161 (2016).

5. A. Pathak, S. Kumar, Biophysical regulation of tumor cell invasion: moving beyond matrix stiffness. Integrative Biology 3, 267–278(2011).

6. A. Pathak, S. Kumar, Independent regulation of tumor cell migration by matrix stiffness and confinement. Proceedings of the National Academy of Sciences 109, 10334–10339(2012).

7. S. R. Vedula et al., Emerging modes of collective cell migration induced by geometrical constraints. Proceedings of the National Academy of Sciences 109, 12974–12979(2012).

8. W. Xi, S. Sonam, T. B. Saw, B. Ladoux, C. T. Lim, Emergent patterns of collective cell migration under tubular confinement. Nat Commun 8, 1517 (2017).

9. J. A. Mitchel et al., In primary airway epithelial cells, the unjamming transition is distinct from the epithelial-to-mesenchymal transition. Nat Commun 11, 5053 (2020).

10. L. Atia et al., Geometric constraints during epithelial jamming. Nature Physics 14, 613–620(2018).

11. J. A. Park et al., Unjamming and cell shape in the asthmatic airway epithelium. Nat Mater 14, 1040–1048(2015).

12. A. Pathak, Scattering of Cell Clusters in Confinement. Biophysical Journal 111, 1496–1506(2016).

13. S. Nasrollahi, A. Pathak, Topographic confinement of epithelial clusters induces epithelial-to-mesenchymal transition in compliant matrices. Sci Rep 6, 18831 (2016).

14. S. Nasrollahi, A. Pathak, Hydrogel-based microchannels to measure confinement- and stiffness-sensitive Yes-associated-protein activity in epithelial clusters. MRS Communications 7, 450–457(2017).

15. P. Friedl, D. Gilmour, Collective cell migration in morphogenesis, regeneration and cancer. Nature Reviews Molecular Cell Biology 10, 445–457 (2009).

16. A. Haeger, K. Wolf, M. M. Zegers, P. Friedl, Collective cell migration: Guidance principles and hierarchies. Trends in Cell Biology 25, 556–566 (2015).

17. A. C. Brown, V. F. Fiore, T. A. Sulchek, T. H. Barker, Physical and chemical microenvironmental cues orthogonally control the degree and duration of fibrosis-associated epithelial-to-mesenchymal transitions. The Journal of Pathology 229, 25–35 (2013).

18. S. C. Wei et al., Matrix stiffness drives epithelial–mesenchymal transition and tumour metastasis through a TWIST1–G3BP2 mechanotransduction pathway. Nature Cell Biology 17, 678–688 (2015).

19. J. S. Park et al., Switch-like enhancement of epithelial-mesenchymal transition by YAP through feedback regulation of WT1 and Rho-family GTPases. Nature Communications 10, 1–15 (2019).

20. J. Lu et al., 14-3-3[zeta] Cooperates with ErbB2 to Promote Ductal Carcinoma In Situ Progression to Invasive Breast Cancer by Inducing Epithelial-Mesenchymal Transition. Cancer Cell 16, 195–207(2009).

21. A. Pathak, S. Kumar, Transforming potential and matrix stiffness co-regulate confinement sensitivity of tumor cell migration. Integrative Biology 5, 1067–1075(2013).

22. S. Nasrollahi, A. Pathak, Topographic confinement of epithelial clusters induces epithelial-to-mesenchymal transition in compliant matrices. Scientific Reports 6, 1–12 (2016).

23. S. Nasrollahi, A. Pathak, Hydrogel-based microchannels to measure confinement- and stiffnesssensitive Yes-associated-protein activity in epithelial clusters. MRS Communications 7, 450–457 (2017).

24. B. Zhao, Q.-Y. Lei, K.-L. Guan, The Hippo–YAP pathway: new connections between regulation of organ size and cancer. Current Opinion in Cell Biology 20, 638–646 (2008).

25. M. Aragona et al., A Mechanical Checkpoint Controls Multicellular Growth through YAP/TAZ Regulation by Actin-Processing Factors. Cell 154, 1047–1059 (2013).

26. T. E. Angelini et al., Glass-like dynamics of collective cell migration. Proceedings of the National Academy of Sciences 108, 4714–4719 (2011).

27. D. Bi, X. Yang, M. C. Marchetti, M. L. Manning, Motility-Driven Glass and Jamming Transitions in Biological Tissues. Physical Review X 6, 021011 (2016).

28. M. Gamboa Castro, S. E. Leggett, I. Y. Wong, Clustering and jamming in epithelial–mesenchymal co-cultures. Soft Matter 12, 8327–8337 (2016).

29. S. Garcia et al., Physics of active jamming during collective cellular motion in a monolayer. Proceedings of the National Academy of Sciences 112, 15314–15319 (2015).

30. J.-A. Park et al., Unjamming and cell shape in the asthmatic airway epithelium. Nature Materials 14, 1040–1048 (2015).

31. L. Atia et al., Geometric constraints during epithelial jamming. Nature Physics 14, 613–620 (2018).

32. J. A. Mitchel et al., In primary airway epithelial cells, the unjamming transition is distinct from the epithelial-to-mesenchymal transition. Nature Communications 11, 1–14 (2020).

33. M. De Marzio et al., Genomic signatures of the unjamming transition in compressed human bronchial epithelial cells. Science Advances 7, eabf1088.

34. D. Bi, J. H. Lopez, J. M. Schwarz, M. L. Manning, A density-independent rigidity transition in biological tissues. Nature Physics 11, 1074–1079 (2015).

35. M. M. Zegers, P. Friedl, Rho GTPases in collective cell migration. Small GTPases 5, e983869–e983869 (2014).

36. S. J. Terry et al., Spatially restricted activation of RhoA signalling at epithelial junctions by p114RhoGEF drives junction formation and morphogenesis. Nature Cell Biology 13, 159–166 (2011).

37. M. Reffay et al., Interplay of RhoA and mechanical forces in collective cell migration driven by leader cells. Nature Cell Biology 16, 217–223 (2014).

38. P. Friedl, K. Wolf, M. M. Zegers, Rho-directed forces in collective migration. Nature Cell Biology 16, 208–210 (2014).

39. J. Ju et al., Optical regulation of endogenous RhoA reveals selection of cellular responses by signal amplitude. Cell Reports 40 (2022).

40. G. Charras, E. Sahai, Physical influences of the extracellular environment on cell migration. Nature Reviews Molecular Cell Biology 15, 813–824 (2014).

41. P. Pandya, J. L. Orgaz, V. Sanz-Moreno, Actomyosin contractility and collective migration: may the force be with you. Current Opinion in Cell Biology 48, 87–96 (2017).

42. C. D. Paul, P. Mistriotis, K. Konstantopoulos, Cancer cell motility: lessons from migration in confined spaces. Nat Rev Cancer 17, 131–140(2017).

43. E. M. Balzer et al., Physical confinement alters tumor cell adhesion and migration phenotypes. FASEB journal : official publication of the Federation of American Societies for Experimental Biology 26, 4045–4056(2012).

44. B. Burkel et al., Preparation of 3D collagen gels and microchannels for the study of 3D interactions In Vivo. Journal of Visualized Experiments 2016, 1–7 (2016).

45. W. Xi, S. Sonam, T. Beng Saw, B. Ladoux, C. Teck Lim, Emergent patterns of collective cell migration under tubular confinement. Nature Communications 8 (2017).

46. S. R. K. Vedula et al., Emerging modes of collective cell migration induced by geometrical constraints. Proceedings of the National Academy of Sciences of the United States of America 109, 12974–12979 (2012).

47. A. K. Marel et al., Flow and diffusion in channel-guided cell migration. Biophysical Journal 107, 1054–1064 (2014).

48. V. Petrolli et al., Confinement-Induced Transition between Wavelike Collective Cell Migration Modes. Physical Review Letters 122, 168101 (2019).

49. S. Varadarajan et al., Mechanosensitive calcium flashes promote sustained RhoA activation during tight junction remodeling. Journal of Cell Biology 221, e202105107 (2022).

50. B. D. Souza, J. Taylor-Papadimitriou (1994) Overexpression of ERBB2 in human mammary epithelial cells signals inhibition of transcription of the E-cadherin gene. in Proc. Nati. Acad. Sci. USA, pp 7202–7206.

51. J. Lu et al., 14-3-3ζ Cooperates with ErbB2 to Promote Ductal Carcinoma In Situ Progression to Invasive Breast Cancer by Inducing Epithelial-Mesenchymal Transition. Cancer Cell 16, 195–207 (2009).

52. A. Pathak, S. Kumar, Transforming potential and matrix stiffness co-regulate confinement sensitivity of tumor cell migration. Integrative Biology 5, 1067–1075 (2013).

53. S.-Z. Lin, W.-Y. Zhang, D. Bi, B. Li, X.-Q. Feng, Energetics of mesoscale cell turbulence in two-dimensional monolayers. Communications Physics 4, 21 (2021).

54. G. Amable et al., Metformin inhibition of colorectal cancer cell migration is associated with rebuilt adherens junctions and FAK downregulation. Journal of Cellular Physiology 235, 8334–8344 (2020).

55. P. Recho, J. Ranft, P. Marcq, One-dimensional collective migration of a proliferating cell monolayer. Soft Matter 12, 2381–2391 (2016).

56. J. J. Fredberg, Power Steering, Power Brakes, and Jamming: Evolution of Collective Cell-Cell Interactions. Physiology 29, 218–219 (2014).

57. M. Abercrombie, J. E. Heaysman, Observations on the social behaviour of cells in tissue culture. I. Speed of movement of chick heart fibroblasts in relation to their mutual contacts. Exp Cell Res 5, 111–131 (1953).

58. T. E. Angelini et al., Glass-like dynamics of collective cell migration. Proceedings of the National Academy of Sciences 108, 4714–4719(2011).

59. A. J. Liu, S. R. Nagel, Jamming is not just cool any more. Nature 396, 21–22 (1998).

60. M. A. Heinrich, R. Alert, A. E. Wolf, A. Košmrlj, D. J. Cohen, Self-assembly of tessellated tissue sheets by expansion and collision. Nature Communications 13, 4026–4026 (2022).

61. M. Reffay et al., Interplay of RhoA and mechanical forces in collective cell migration driven by leader cells. Nature Cell Biology 16, 217–223(2014).

62. E. Anear, R. W. Parish, The effects of modifying RhoA and Rac1 activities on heterotypic contact inhibition of locomotion. FEBS Letters 586, 1330–1335 (2012).

63. J. Schindelin et al., Fiji: an open-source platform for biological-image analysis. Nature Methods 9, 676–682 (2012).

64. D. Ershov et al., TrackMate 7: integrating state-of-the-art segmentation algorithms into tracking pipelines. Nature Methods 19, 829–832 (2022).

65. D. N. Mashburn, H. E. Lynch, X. Ma, M. S. Hutson, Enabling user-guided segmentation and tracking of surface-labeled cells in time-lapse image sets of living tissues. Cytometry Part A 81A, 409–418 (2012).

